# Quorum sensing N-Acyl homoserine lactones are a new class of anti-schistosomal

**DOI:** 10.1101/2020.07.24.219311

**Authors:** H Whiteland, A Crusco, LW Bloemberg, J Tibble-Howlings, J Forde-Thomas, A Coghlan, P. J Murphy, KF Hoffmann

## Abstract

**Background:** Schistosomiasis is a prevalent neglected tropical disease that affects approximately 300 million people worldwide. Its treatment is through a single class chemotherapy, praziquantel. Concerns surrounding the emergence of praziquantel insensitivity have led to a need for developing novel anthelmintics.

**Methodology/Principle findings:** Through evaluating and screening fourteen compounds (initially developed for anti-cancer and anti-viral projects) against *Schistosoma mansoni*, one of three species responsible for most cases of human schistosomiasis, a racemic N-acyl homoserine (**1**) demonstrated good efficacy against all intra mammalian lifecycle stages including schistosomula (EC_50_ = 4.7 µM), juvenile worms (EC_50_ = 4.3 µM) and adult worms (EC_50_ = 8.3 µM). To begin exploring structural activity relationships, a further 8 analogues of this compound were generated, including individual (*R*)- and (*S*)- enantiomers. Upon anti-schistosomal screening of these analogues, the (*R*)- enantiomer retained activity, whereas the (*S*)- lost activity. Furthermore, modification of the lactone ring to a thiolactone ring (**3**) improved potency against schistosomula (EC_50_ = 2.1 µM), juvenile worms (EC_50_ = 0.5 µM) and adult worms (EC_50_ = 4.8 µM). As the active racemic parent compound is structurally similar to quorum sensing signaling peptides used by bacteria, further evaluation of its effect (along with its stereoisomers and the thiolactone analogues) against Gram^+^ (*Staphylococcus aureus*) and Gram^-^ (*Escherichia coli*) species was conducted. While some activity was observed against both Gram^+^ and Gram^-^ bacteria species for the racemic compound **1** (MIC 125 mg/L), the (*R*) stereoisomer had better activity (125 mg/L) than the (*S*) (>125mg/L). However, the greatest antimicrobial activity (MIC 31.25 mg/L against *S. aureus*) was observed for the thiolactone containing analogue (**3**).

**Conclusion/Significance:** To the best of our knowledge, this is the first demonstration that N-Acyl homoserines exhibit anthelmintic activities. Furthermore, their additional action on Gram^+^ bacteria opens a new avenue for exploring these molecules more broadly as part of future anti-infective initiatives.

**Author Summary:** Schistosomiasis, caused by infection with blood fluke schistosomes, is a neglected tropical disease that negatively impacts the lives of approximately 300 million people worldwide. In the absence of a vaccine, it is currently controlled by a single drug, Praziquantel (PZQ). Although incredibly valuable in controlling disease burden, PZQ-mediated chemotherapy is ineffective against juvenile worms and may not be sustainable should resistance develop. The need to identify an alternative or combinatorial drug is, therefore, a priority in contributing to the control of this parasitic disease into the 21^st^ century. In this study, we have identified a new class of anthelmintic, N-acyl homoserine lactones, which are normally used by bacteria for quorum sensing and population density control. The tested N-acyl homoserine lactones were active against all intra-human schistosome lifecycle stages, in particular, when a thiolactone modification to the core N-acyl homoserine ring was made. Interestingly, these N-acyl homoserine lactones also displayed antimicrobial activities against Gram^+^ *Staphylococcus aureus*. By demonstrating broad activities against schistosomes and bacteria exemplars, this study identified a potential route for the further development of a new anti-infective class.

## Introduction

The development of drug resistant prokaryotic and eukaryotic pathogens is of great concern for the sustainable control of both human and animal diseases; therefore, the need for new anti-infectives is a global health priority. One notorious group of difficult to control pathogens are those that cause the Neglected Tropical Diseases (NTDs). In total, 17 parasites/microbes and their related infections are now identified as responsible for most NTDs worldwide [1], with schistosomiasis, leishmaniasis and soil-transmitted helminthiasis causing significant disability adjusted life years lost per annum [2]. As vaccines are unavailable for the prevention of most NTDs, a small number of chemotherapies remain the primary means of global control. However, drug resistance or reduced susceptibility to these limited drug classes has been reported for NTD-causing bacteria [3], fungi [4], helminths [5, 6] and protozoa [7, 8]. These chemotherapy limitations have prompted an urgent need for research into the development of new anti-infective agents. Nevertheless, funding to support this agenda is unlikely to originate from the pharmaceutical sector due to the lack of financial returns associated with controlling diseases predominantly affecting low to middle-income countries [9-11]. Thus, philanthropic and public organisations are often driving the majority of new anti-infective initiatives targeting the NTDs with derived funding supporting research conducted in higher education or research institutes.

One particular debilitating NTD, schistosomiasis, affects over 300 million people worldwide [12] and is predominantly caused by infection with three *Schistosoma* species [13]. Therapeutic treatment involves praziquantel (PZQ) as the frontline control strategy. However, PZQ is ineffective against the juvenile stage of the parasite *in vivo*, which necessitates repeat administrations to reach maximal efficacy in endemic populations [14]. Furthermore, PZQ is currently produced as a racemic mixture and only the (*R*) - enantiomer is active; the (*S*) - enantiomer contributes to some of the side effects including bitter taste and non-compliance in the young [15]. These drug-related limitations, together with a constant fear of PZQ resistance developing, has fuelled investigations into the identification of PZQ replacement or combinatorial anti-schistosomal drugs.

Towards this goal, our group has recently identified several diverse starting points for anti-schistosomal drug discovery. These include diterpenoids [16, 17], triterpenoids [18], and epigenetic probes/inhibitors [19, 20]. While the primary focus of these investigations evaluated compound-induced activity on *Schistosoma mansoni* (the schistosome species responsible for both Old and New world schistosomiasis), parallel studies were also conducted to quantify anti-infective activities against other NTD-causing pathogens including *Fasciola hepatica* (liver fluke) [16-18] or NTD models such as *Mycobacterium smegmatis* (related to *Mycobacterium leprae*) [21].

In this present study, we assessed the anti-schistosomal activity of 14 in house prepared synthetic compounds (or intermediate analogues of these compounds). As the most active anti-schistosomal compounds were structurally similar to N-acyl homoserines, a class of signalling molecule involved in bacterial quorum sensing and population density control [22], we additionally investigated their anti-microbial activity. Amongst the compounds tested, one demonstrated moderate activity against Gram^+^ (*Staphylococcus aureu*s) bacteria. Our collective results demonstrate that N-acyl homoserines represent a new class of anthelmintics with additional activity against bacteria. Further development of these molecules could be pursued as promising new chemotherapeutics for schistosomiasis and other NTDs.

## Materials and Methods

### Ethics statement

All procedures performed on mice (project licenses 40/3700 and P3B8C46FD) adhered to the United Kingdom Home Office Animals (Scientific Procedures) Act of 1986 as well as the European Union Animals Directive 2010/63/EU and were approved by Aberystwyth University’s (AU) Animal Welfare and Ethical Review Bodies (AWERB).

### Synthetic Methodology

Lactones (compounds **1-9)** were prepared by the reaction of the lactone/thiolactone, with the required acyl halide in the presence of NEt_3_ in chloroform or potassium carbonate in a two phase water/chloroform mix (Scheme 1). Yields and conditions are shown in Table 1 and full synthetic and spectroscopic details are found in S1 Protocol.

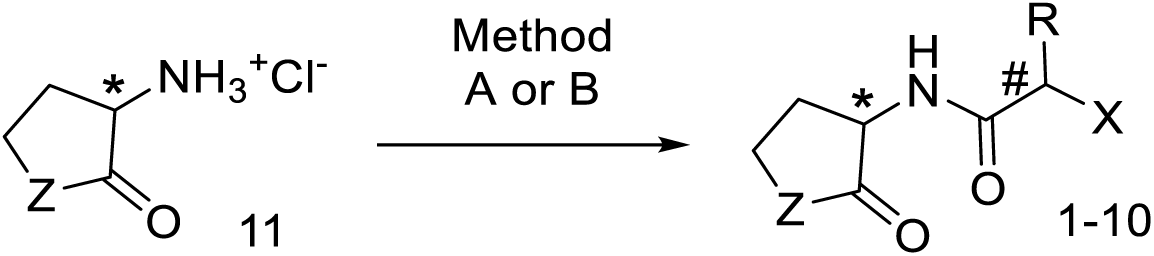
Scheme 1: Method A: NEt_3_, CHCl_3_, 0°C; Method B: K_2_CO_3_, CH_2_Cl_2_/H_2_O, 0°C. R = H. Me, X = O, S.* Chiral centres

**Table 1:** Preparatory summary of compounds 1-9.

| Compound | X | Z | R | * | # | Method | Yield | Mp/°C |
| --- | --- | --- | --- | --- | --- | --- | --- | --- |
| **1** | Br | O | H | Racemic | NA | A | 75% | 95-97 |
| **2** | Br | O | Me | Racemic | Racemic | A | 33% | 157-69 |
| **3** | Br | S | H | Racemic | NA | B | 40% | 111-3 |
| **4** | Br | S | Me | Racemic | Racemic | B | 73% | 125-28 |
| **5** | Br | O | H | *R* | NA | B | 25% | 130-3 |
| **6** | Br | O | H | *S* | NA | B | 14% | 130-3 |
| **7** | Cl | O | H | Racemic | NA | A | 68% | 111-5 |
| **8** | Cl | S | H | Racemic | NA | B | 61% | 124-7 |
| **9** | Cl | O | Me | Racemic | Racemic | A | 65% | 148-50 |

The symbols “\*” and “#” refer to the absolute configuration at the carbon centres indicated in scheme 1.
NA= Not applicable
Z = O, S, R = H, Me.

Scheme 1: Method A: NEt_3_, CHCl_3_, 0°C; Method B: K_2_CO_3_, CH_2_Cl_2_/H_2_O, 0°C. R = H. Me, X = O, S.* Chiral centres

### Compound storage and handling

All compounds were solubilised in DMSO (Fisher Scientific, UK) to a stock concentration of 10 mM and stored at -20°C until required. Positive controls for *S. mansoni* screens included PZQ (Sigma-Aldrich, UK) and auranofin (Sigma-Aldrich, UK), which were also treated in the same manner as the test compounds.

### Screening of *S. mansoni* schistosomula

*Biomphalaria glabrata* (NMRI and the previously described pigmented strains [23]) snails infected with *S. mansoni* (Puerto Rican strain) were shed for 2 hrs under light at 26 °C. Cercariae were collected, mechanically transformed into schistosomula [24] and subsequently prepared for high throughput screening (HTS) on the Roboworm platform as previously described [25]. Compounds were initially tested at a final concentration of 10 μM and those that were active were further titrated at concentrations of 10, 5, 2.5, 1.25 and 0.625 µM. EC_50_ values were calculated from the titrated concentrations by non-linear regression, after log transformation of concentrations and data normalization using GraphPad Prism 7.02.

### Screening of adult *S. mansoni* blood flukes (7-week worms)

Adult *S. mansoni* parasites were recovered by hepatic portal vein perfusion from TO mice (Harlan, UK) that were percutaneously infected seven weeks earlier with 180 cercariae. Three adult worm pairs per well, in duplicate, were transferred into 48 well plates (Fisher Scientific, Loughborough, UK) and cultured at 37 °C in an atmosphere containing 5% CO_2_ in DMEM (Gibco, Paisley, UK) containing 10% v/v HEPES, 10% v/v Foetal Bovine Serum (FBS), 0.7% v/v 200 mM L-Glutamine and 1X v/v penicillin-streptomycin. Worms were dosed with test compounds at 20 µM, 10 µM, 5 µM, 2.5 µM, 1.25 µM and 0.625 µM (in 0.2% DMSO) for 72 hr. Adult worms were scored manually at 72 hr using the WHO-TDR metric scoring system as described previously [26]. Dose response curves and EC_50_ values were obtained by non-linear regression, after log transformation of concentrations and data normalization using GraphPad Prism 7.02. At 72 hr, the medium from each well was also collected, centrifuged at 1000 rpm for 2 min. Afterwards, the supernatant was removed, and the remaining egg pellet re-suspended in 10% v/v formalin. Eggs that were oval and contained a fully formed lateral spine were subsequently counted.

### Screening of juvenile *S. mansoni* blood flukes (3-week worms)

Juvenile *S. mansoni* parasites were recovered via hepatic portal vein perfusion from TO mice (Harlan, UK) that were infected percutaneously three weeks earlier with 4000 cercariae. Preparation and centrifugation of juvenile worms have been described previously [18]. Briefly, juvenile worms (n=13-33 individuals/well) in 200 µl of a 96-well tissue culture plate were co-cultured with compounds (15 µM, 7.5 µM, 3.75 µM, 1.83 µM, 0.94 µM and 0.47 µM (in 1.25% DMSO) in DMEM (Gibco, Paisley, UK) supplemented with 10% v/v Hepes (Sigma-Aldrich, Gillingham, UK), 10% v/v FBS (Gibco, Paisley, UK), 0.7 % v/v 200 mM L-Glutamine (Gibco, Paisley, UK) and 1X v/v penicillin-streptomycin (Fisher Scientific, UK). Positive control wells included either PZQ or auranofin (15 µM in 1.25% DMSO) whereas negative wells included DMSO (1.25%). Parasites were incubated at 37 °C in an atmosphere containing 5% CO_2_ for 72 hr at which time worm motility was scored between 0 and 4: 0 = dead, 1 = movement of the suckers only and slight contraction of the body, 2 = movement at the anterior and posterior regions only, 3 = full body movement but sluggish and 4 = normal movement. After motility quantification, 2 µg/mL of PI was added to each well and the plate returned to 37°C, 5% CO_2_ for 15 minutes [27]. Each well was subsequently imaged on the Roboworm platform using brightfield and fluorescent microscopy (excitation wavelength = 580 nm; emission wavelength = 604 nm). The number of PI positive vs PI negative juvenile worms were cross-checked with the motility scores obtained by our scoring matrix, and the data reported as percentage of PI positive across all parasites within the well. EC_50_ values were calculated from the motility scores obtained from the dose response titration (as detailed above) and dose response curves were obtained by non-linear regression, after log transformation of concentrations and data normalization using GraphPad Prism 7.02.

### Cell Cytotoxicity Assays

The cytotoxicity of each compound was assessed on human HepG2 cells as described previously [25]. Briefly, 2 × 10^4^ cells/well were seeded in black walled 96-well microtiter plates (Fisher Scientific, Loughborough, UK) and incubated for 24 hr at 37°C in a humidified atmosphere containing 5% CO_2_. To each well, compounds were subsequently added to obtain final concentrations (in 1% DMSO) of 100 µM, 75 µM, 50 µM, 25 µM, 10 µM and 5 µM. Following a further incubation for 24 hr, the MTT assay was performed as previously described [25, 28]. Dose response curves were obtained by non-linear regression, after log transformation of concentrations and data normalization using GraphPad Prism 7.02.

### Bacterial Growth, Minimum Inhibitory Concentration (MIC) Calculation, and EC_50_ Determination

*S. aureus* ATCC 29213 and *E. coli* ATCC 25922 were cultured in Luria-Bertani (LB) medium at 37 °C with aeration at 200 rpm for 24 hr, with all procedures performed in a biosafety level 2 (BSL2) cabinet. Stationary phase cultures were then used for minimum inhibitory concentration (MIC) determination using the broth microdilution method, in fresh LB medium, in a 96-well plate [29]. All compounds were tested in triplicate using an initial bacterial concentration of 5.0 × 10^5^ colony forming units (CFU)/mL at a final concentration of 125 mg/L (5% and 2.5% v/v methanol). Compounds with no visible growth at 125 mg/L were further evaluated with progressing dilutions. The MIC was determined as the lowest concentration of a compound at which no growth was visible after 24 hr. Dilutions were repeated in three independent experiments where the optical density (OD_600_) was measured in a Hidex plate spectrophotometer and absorbance data used for the calculation of an IC_50_ value. The EC_50_ value was obtained from a dose response titration (125–0.09 mg/L). Dose response curves were obtained by non-linear regression, after log transformation of concentrations and data normalization using GraphPad Prism 7.02.

### Bioinformatics

The names and structures (SMILES strings) of chemical compounds were identified in PubMed abstracts in June 2019 using the chemistry text-mining software LeadMine v 3.1.2 (NextMove Software Ltd.) [30]. Chemical compounds were identified in all PubMed abstracts containing any one of long list of words relating to schistosomes and anthelmintic/antiparasitic compounds (e.g. ‘schistosoma’, ‘sporocyst’, ‘sporocysts’, ‘miracidium’, ‘miracidia’, ‘somule’, ‘somules’, ‘schistosomula’, ‘schistosomulum’, ‘cercariae’, ‘cercaria’, ‘schistosome’, ‘schistosomes’, ‘antiparasitic’, ‘antinematodal’, ‘anthelmintic’, etc.). The structures of our screening hits were compared to those of the chemical compounds identified in PubMed abstracts using DataWarrior [31], by using DataWarrior’s Similarity Analysis function with its FragFP descriptor.

A map of the quorum sensing pathway in *Pseudomonas aeruginosa* was found using the Kyoto Encyclopedia of Genes and Genomes (KEGG) database. Through this database, the NCBI Protein IDs and the corresponding amino acid sequences were obtained for LasR (NP_250121), LasI (NP_250123), RhIR (NP_252167), RHII (NP_252166). These sequences were used as queries for a protein BLAST (BLASTp) search against the *S. mansoni* genome in Wormbase-Parasite [32].

### Statistics

All Statistical analyses were conducted using GraphPad Prism 7 software. To determine significant differences amongst population means, a Kruskal-Wallis ANOVA followed by Dunn’s multiple comparisons test was used. *p* values are indicated as follows: * <0.05, ** <0.01, *** <0.001.

## Results

As part of our anti-infective research activities, a total of 14 synthetic compounds (including some intermediate analogues; S1 Table) were entered into a screening pipeline to identify active molecules (Fig. 1).

**Figure 1:**
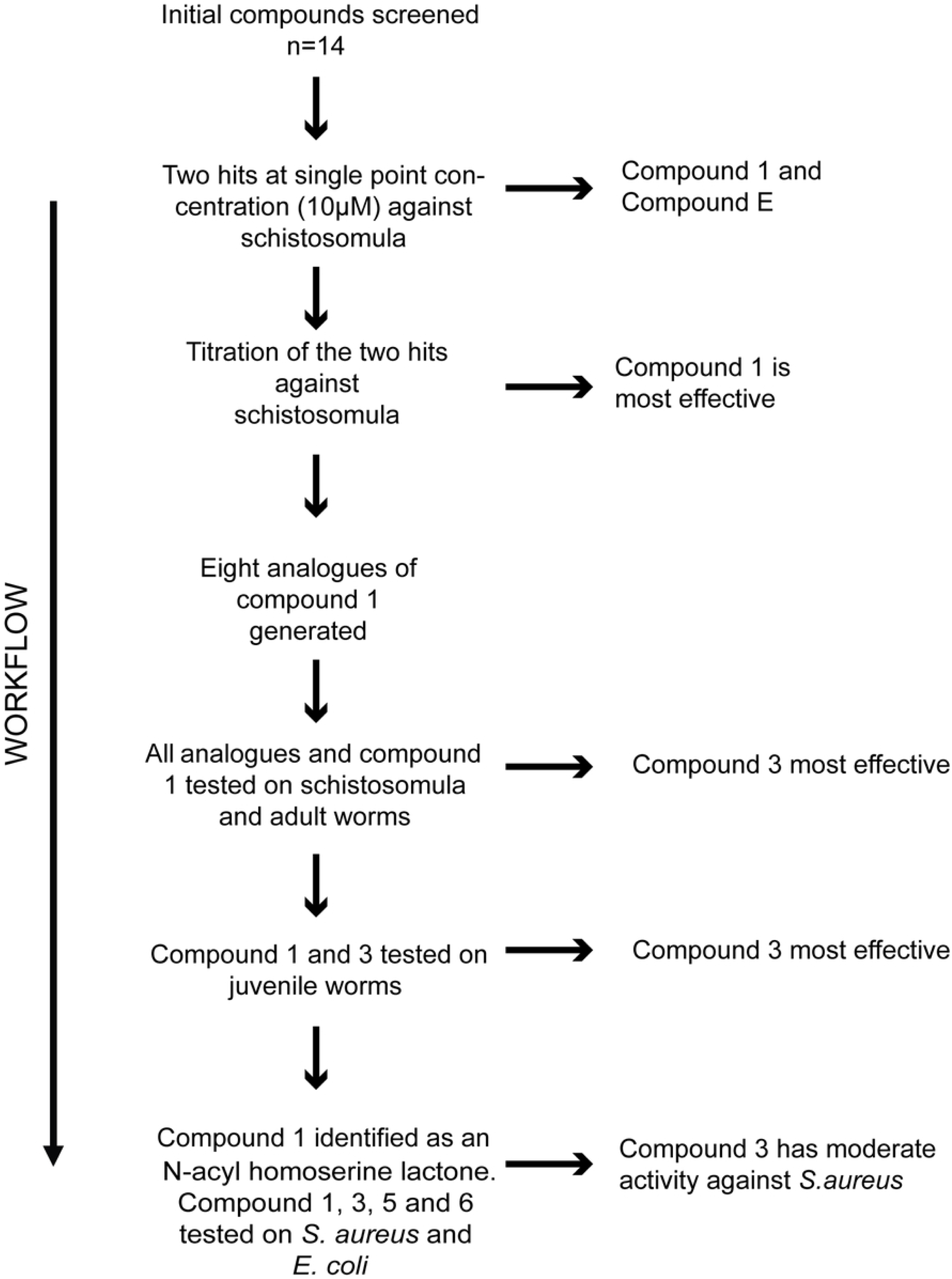
The screening pipeline utilised in this anti-infective study. A total of 14 compounds were initially screened against *S. mansoni* schistosomula at a final concentration of 10 µM. Hits were subsequently subjected to dose response titrations with the most active anti-schistosomula compound (Compound **1**) subsequently being used as a template for the preparation of further derivatives (compounds **2-9**). Compounds **2-9** were subsequently subjected to dose response titrations against schistosomula and adult worms. The most active compounds (original compound **1** and analogue **3**) were next titrated against juvenile worms. Finally, compounds **1, 3, 5** and **6** were additionally titrated against both Gram^+^ (*S. aureus*) and Gram^-^ (*E. coli*) bacterial exemplars.

These 14 compounds (S1 Table) were initially screened against the larval stage of *S. mansoni* (schistosomula) (S1 Figure). Amongst the collection, two compounds negatively affected both schistosomula motility and phenotype metrics at 10 µM (Compound **1** and Compound **E** (S1a Figure)). Subsequent titration of these two compounds demonstrated that Compound **1** was more active (an average EC_50_ of 4.7 µM for phenotype and motility) than Compound **E** (an average EC_50_ of 5.6 µM for phenotype and motility (S1b Figure)). Due to these initial anti-schistosomula screens identifying compound **1** as being moderately potent, a total of 8 analogues, all containing a lactone ring core structure but differing in functional group modifications, were synthesised to further assess anti-schistosomal activities (S1 Table).

Firstly, in a direct comparison to compound **1**, each of the 8 analogues as well as compound **1** were titrated against the schistosomula stage to assess compound-induced changes to parasite motility and/or phenotype (Fig. 2).

**Figure 2:**
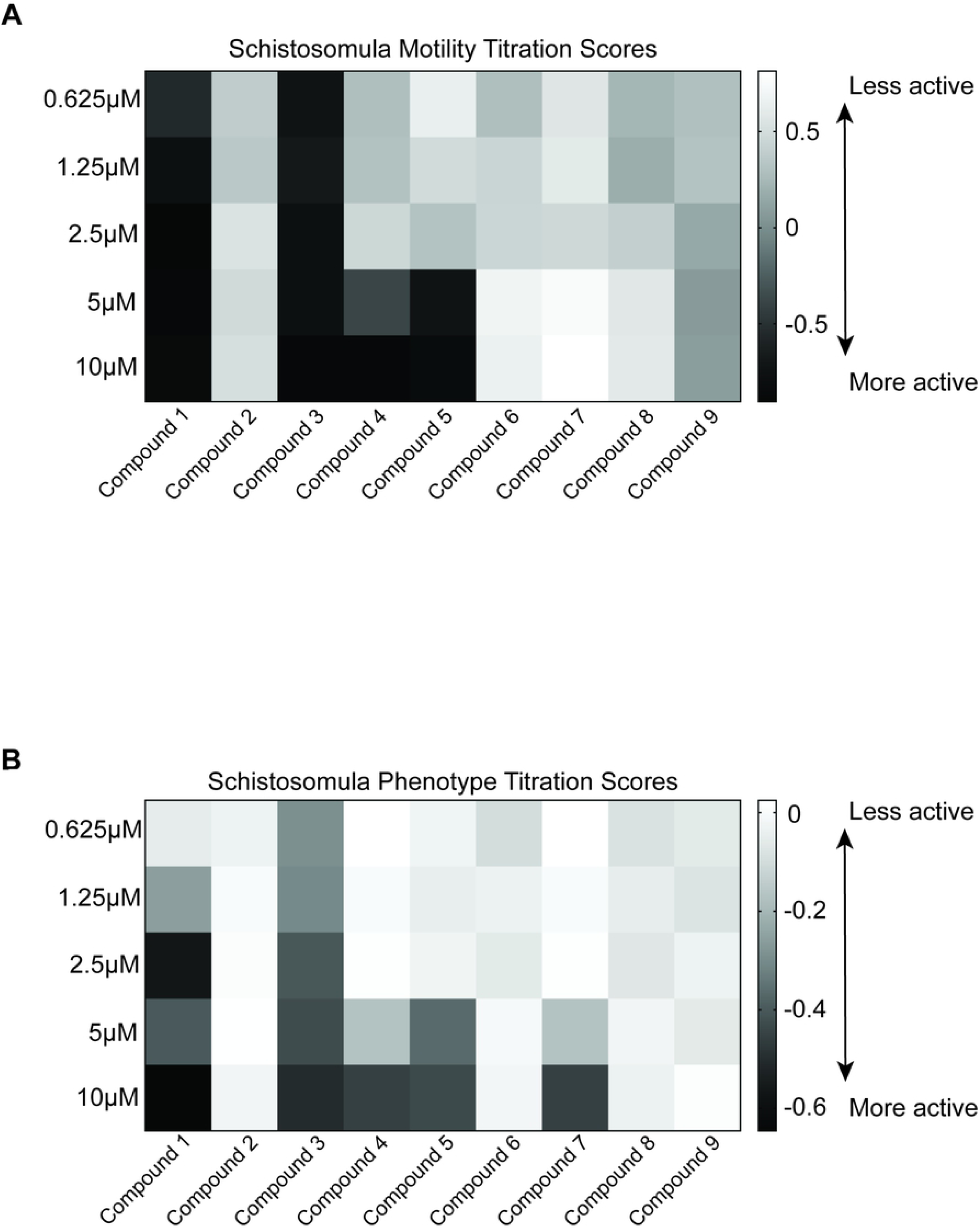
Anti-schistosomula activity of the eight analogues compared to parent compound 1. A total of 120 mechanically transformed schistosomula were co-cultured with each compound, titrated at doses between 10 and 0.625 µM. Test plates were incubated at 37°C for 72 hrs in an atmosphere containing 5% CO_2._ At 72hrs, schistosomula were scored using the Roboworm platform for both motility (**A**) and phenotype (**B**). Any compound that induced a score of below -0.35 for motility (**A**) and -0.15 for phenotype (**B**) were considered a hit. Black squares indicate the most positive effect on motility or phenotype; grey scale from dark grey to lighter shades of grey indicates a progressive reduced compound efficacy; white squares indicate no effect on either phenotype or motility. Z’ values for this screen was 0.41741 for motility and 0.57275 for phenotype.

Out of the nine compounds titrated, four affected schistosomula motility (44.4%; compounds **1, 3, 4** and **5**) (Fig. 2A). Compounds **4** and **5** affected the motility of the parasites at both 10 and 5 µM, whereas compound **1** and **3** affected the motility at all concentrations tested. When evaluating compound-mediated alterations of schistosomula phenotypes, five of the nine compounds had an effect (55.6%; compounds **1, 3, 4, 5** and **7**) (Fig. 2B). Analogues **4, 5** and **7** affected the phenotype of the parasites at 10 and 5 µM, compound **1** affected this metric from 10 to 1.25 µM and compound **3** affected the phenotype of the schistosomula at all concentrations tested. Therefore, a substitution of the simple lactone ring (**1**) for a thiolactone ring (**3**) increased comparative anti-schistosomula potency at all concentrations tested. Of particular interest is the comparison of the two enantiomeric forms of Compound **1**. The (*R*)-enantiomer (**5**) affected schistosomula for both phenotype and motility at 10 µM and 5 µM concentrations. However, no anti-schistosomula activity was observed for the (*S*)-enantiomer (**6**), suggesting that stereo-specificity of N-acyl homoserine is critical to structural activity relationships (SAR).

Due to initial indications that these analogues had varying activities against the schistosomula lifecycle stage (Table 2), the parent compound and its 8 analogues (Compounds **1**-**9**) were subsequently screened against 7-week old adult male and female worms (Fig. 3).

**Table 2:**
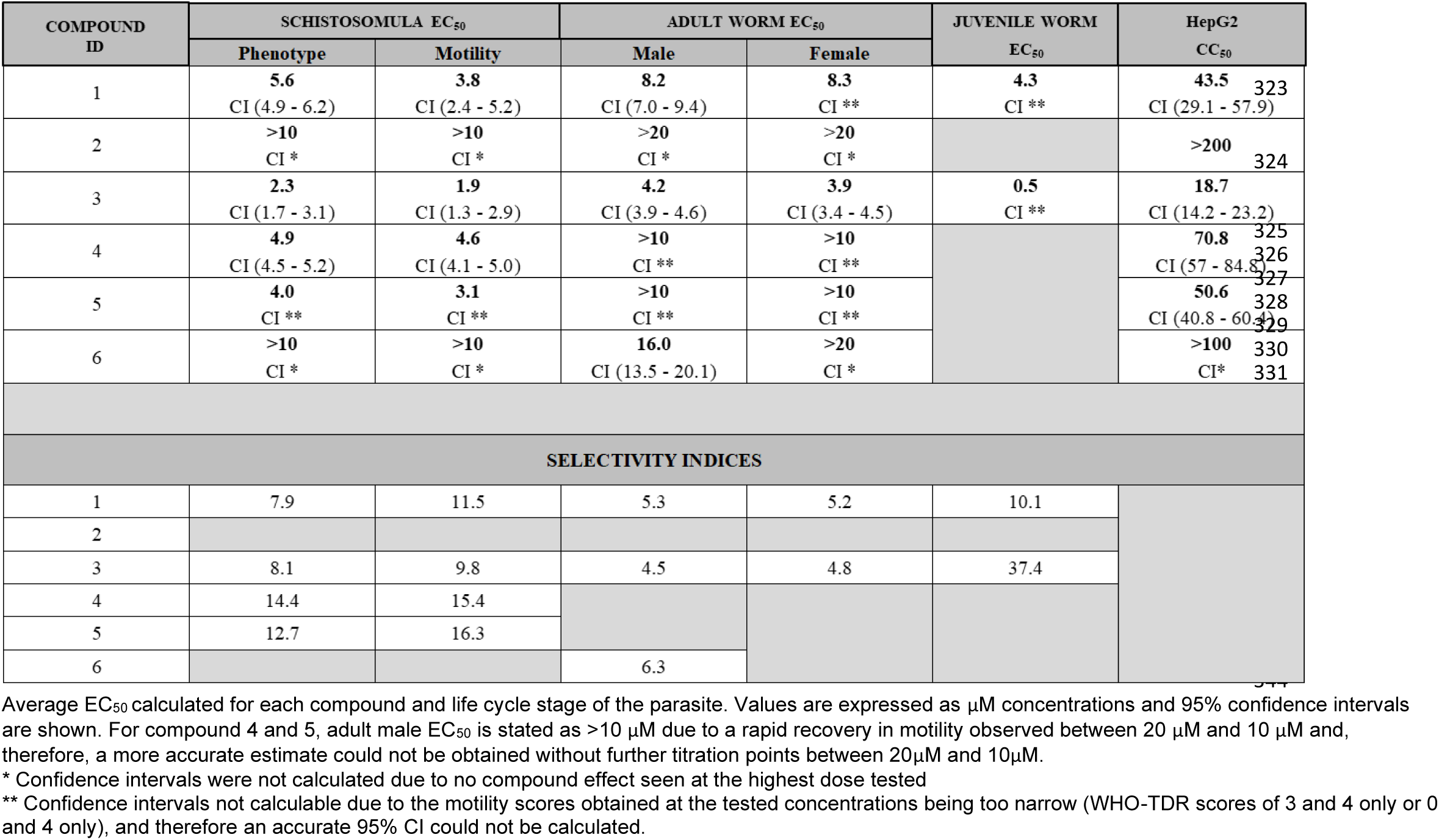
Calculated EC_50,_ CC_50_ and subsequent selectivity indices of N-acyl homoserines.

**Figure 3:**
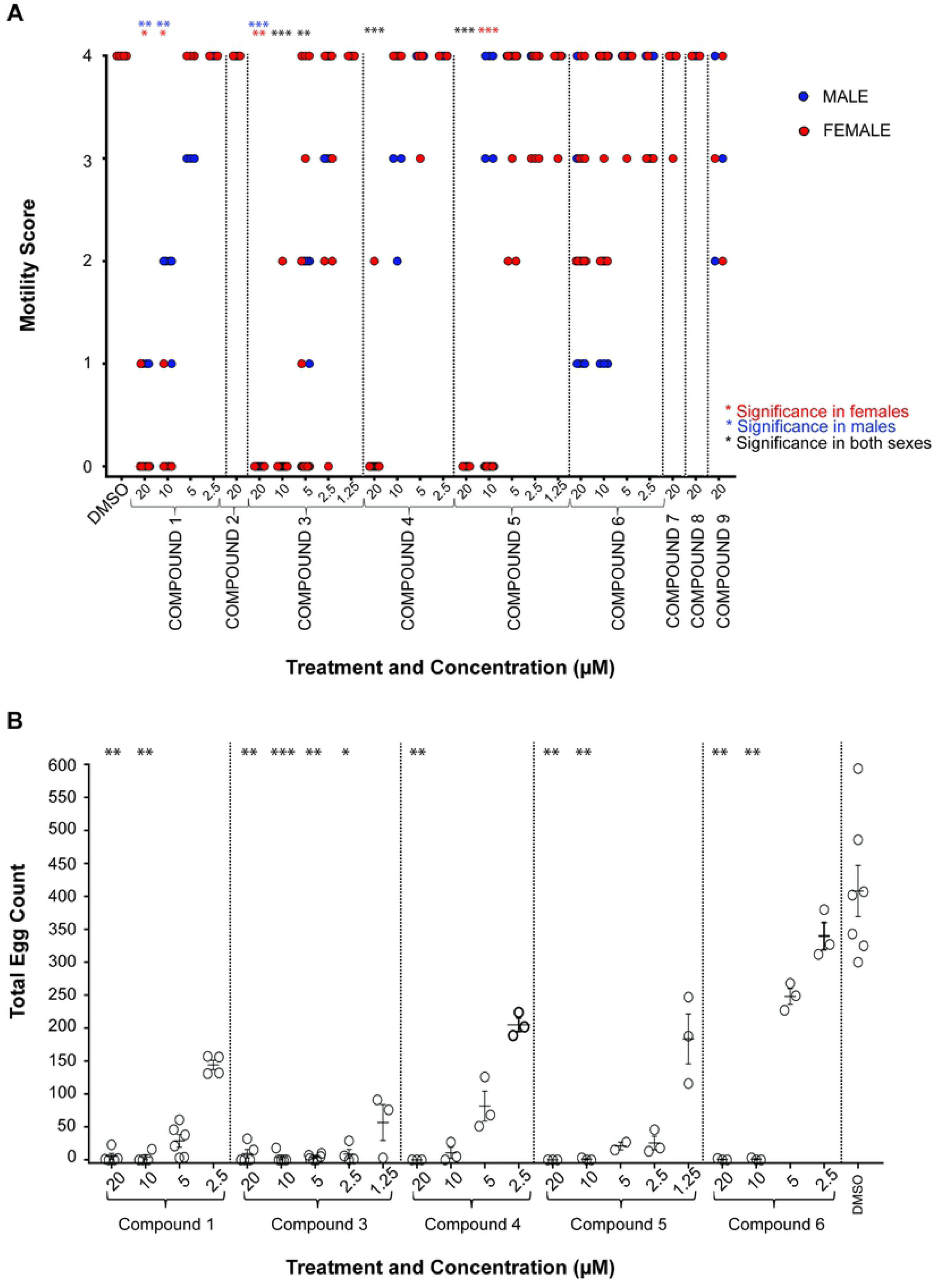
Adult schistosome motility and egg production are differentially affected by the nine N-acyl homoserine lactones. **A**) Adult *S. mansoni* worm pairs were cultured in decreasing compound concentrations of between 20 µM and 1.25 µM (not all concentrations were used for all compounds) for 72 hrs at 37°C, 5% CO_2._ Parasite motility was evaluated for each sex and scored using the WHO-TDR scoring system (0= Dead parasite, 4= Normal/Healthy movement). **B)** Culture media were collected from some adult co-cultures at 72 hrs and eggs present in the media were counted. *p* values are indicated as follows: * <0.05, ** <0.01, *** <0.001.

Initially, these screens were conducted at 20 µM; any compounds that scored a 1 or 0 for motility at this primary concentration were further titrated until no further effect was observed. For Compound **1**, an EC_50_ for males was estimated to be 8.2 µM and for females 8.3 µM (Fig. 3A). Similar to the schistosomula assays, the differences observed in adult parasite motility when treated with the (*R*)- and (*S*)- enantiomers of Compound **1** were striking. The (*R*)-enantiomer (**5**) caused complete immobility at the highest dose of 20 µM; conversely, the (*S*)-enantiomer (**6**) did not have this effect. Estimated EC_50_s for male and female parasites for these two enantiomers were determined (Table 2), with Compound **5** ((*R*)- enantiomer) having greater activity than Compound **6** ((*S*)- enantiomer).

While the propanamide analogue (**2**) had no activity against the adults, the thiolactone analogue (**3**) had good activity (similar to its effects on schistosomula) with significant reduction in motility observed down to 5 µM; EC_50_s of 4.2 µM for males and 3.9 µM for females were determined (Table 2). The thiolactone propanamide analogue (**4**) only demonstrated lethality at the highest concentration of 20 µM with an EC_50_ of 10.5 µM and 18.4 µM for males and females respectively (Table 2). No lethality/motility defects were observed for any of the chloro-analogues (Compounds **7, 8** and **9**).

As morbidity associated with schistosomiasis is caused by egg-induced granuloma formation in tissues and the subsequent development of fibrotic lesions around these granulomas [33], compound-mediated modulation of *in vitro* egg production (as a surrogate for the *in vivo* pathology initiator) was next assessed (Fig. 3B). Specifically, media derived from adult worm cultures incubated with compounds that resulted in complete immobility/lethality (compounds **1, 3, 4**, and **5**; compound **6** was also included as the (*S*)- stereoisomer of compound **1**) were collected and eggs counted (Fig. 3B). In comparison to the negative control DMSO, in which egg counts ranged from 300 – 594 (Average - 408), all N-Acyl homoserine lactones affected fecundity. The parent Compound (**1**) significantly affected egg laying down to 10 µM (*p* < 0.0022) when compared to DMSO. While both (*R*)-(compound **5**) and (*S*)- (compound **6**) enantiomers reduced egg production, the *(R)*- enantiomer was more effective. Of the compounds evaluated, Compound **3** was, once again, the most potent. Here, fecundity was significantly reduced at concentrations down to 2.5 µM (*p* = 0.0413) when compared to DMSO controls. While egg production was still affected at 1.25 µM, this was not statistically significant (*p* > 0.9999).

Next, we further evaluated the most potent analogue (compound **3**) on 3-week old juvenile worms and compared its effect to that induced by the racemic parent compound (**1**) (Fig. 4).

**Figure 4:**
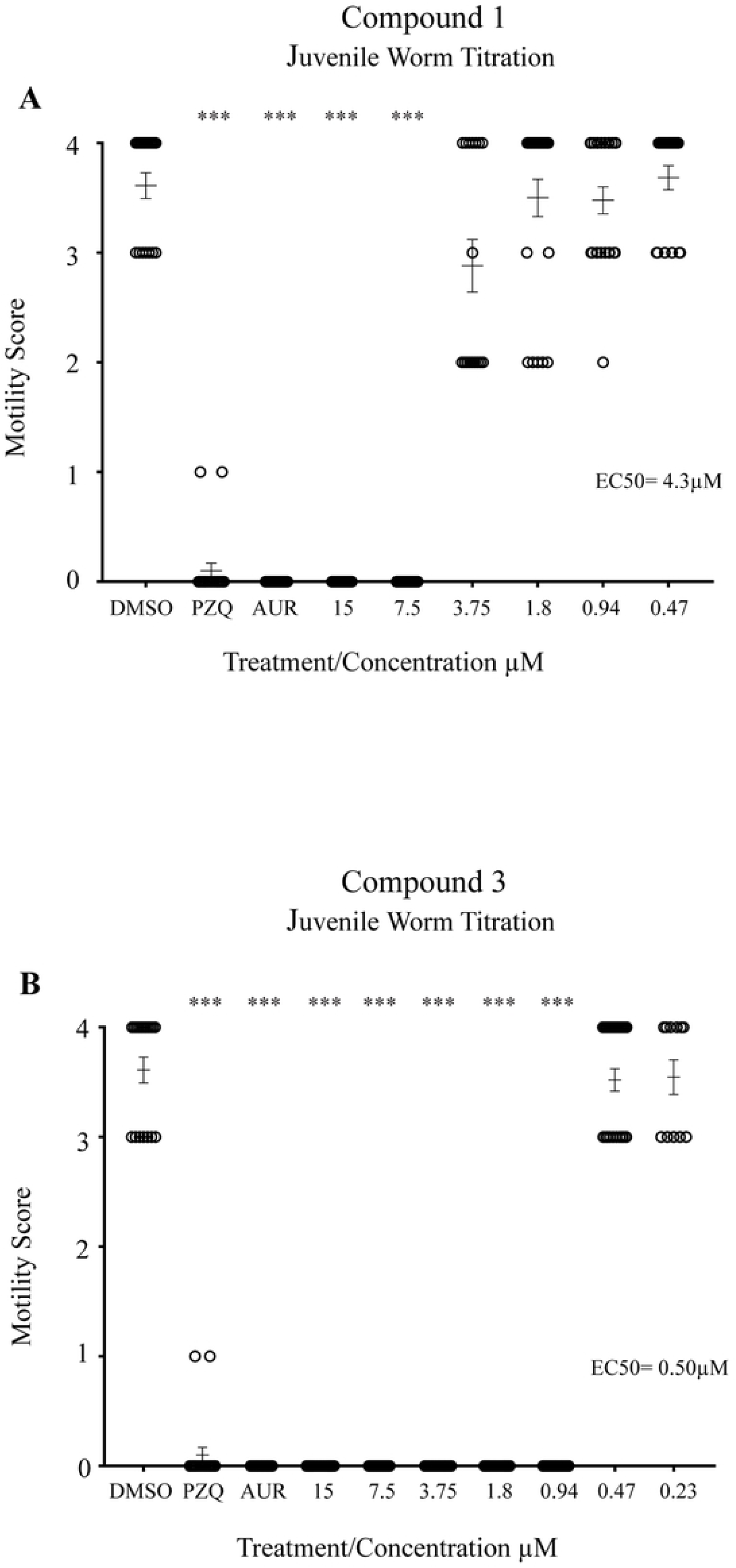
Three week juvenile worms are immobilised by N-Acyl homoserine lactones. Three-week old juvenile *S. mansoni* worms (n= 13-33 per well) were co-cultured with compounds (**1**) and (**3**) at concentrations spanning 15 µM - 0.23 µM for 72 hrs at 37°CC in a humidified atmosphere containing 5% CO_2_. Parasite motility was scored between 0 – 4 (0 = no movement/dead, 4 = full movement/healthy). DMSO negative controls were also included (1.25% final concentration) as well as two positive controls (15 µM PZQ and 15 µM auranofin; both in 1.25% DMSO). **A)** Effect of compound (**1**) on juvenile worm motility. **(B)** Effect of compound (**3**) on juvenile worm motility. *p* values are indicated as follows: * <0.05, ** <0.01, *** <0.001.

For compound **1**, an EC_50_ of 4.3µM was noted with significant effect on worm motility down to 7.5 µM (*p* = <0.0001) (Fig. 4A). For the thiolactone analogue (**3**), complete immobility was observed down to 0.94 µM (*p* <0.0001 for all concentrations) and an EC_50_ of 0.5 µM determined (Fig 4B). This data is consistent with the findings observed in schistosomula and the 7-week adult worm screens where incorporation of a thiolactone resulted in more effective anti-schistosomal activity. To additionally demonstrate that these N-Acyl homoserine lactones led to juvenile worm death, propidium iodide was utilised in compound **1** and **3** co-cultures (Fig. 5).

**Figure 5:**
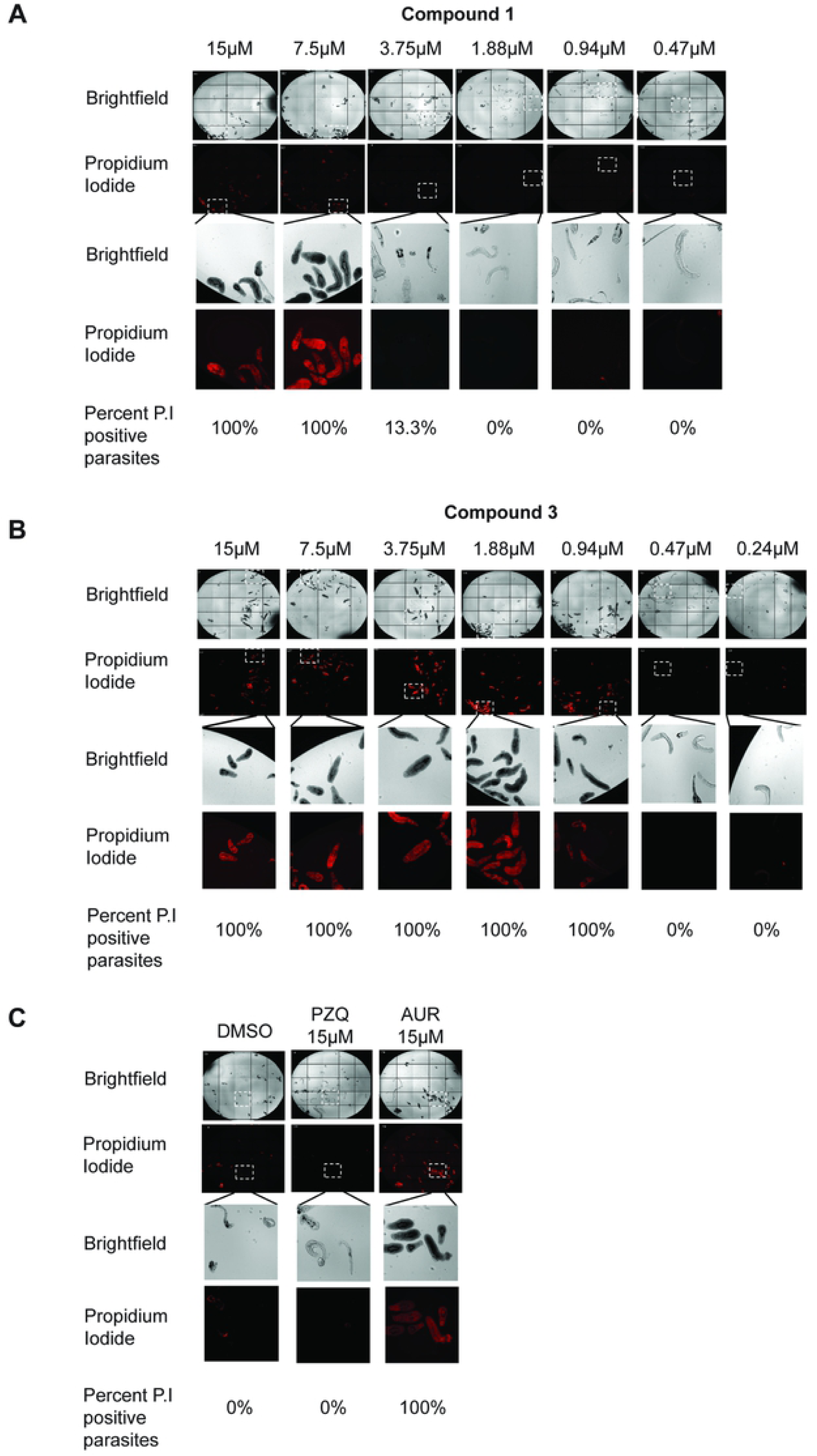
N-Acyl homoserine lactones kill juvenile stage schistosomes. Three week juvenile schistosomes, incubated with compounds (**1**), (**3**) or controls for 72 hrs, were subsequently cultured with PI at a final concentration of 2 µg/mL for 15 minutes at 37 °C in an environment containing 5% CO_2_. PI positive parasites (dead) were counted and the percent live vs dead in each well is indicated. **A)** Quantification of compound (**1**) mediated juvenile death. **B)** Quantification of compound (**3**) mediated juvenile death. **C)** Quantification of juvenile deaths in co-cultures containing 15 µM PZQ (in 1.25% DMSO), 15 µM auranofin (in 1.25% DMSO) and 1.25% DMSO.

The number of PI positive parasites when dosed with compound **1** was 100% at 15 and 7.5 µM concentrations, 13.3% at 3.75 µM and 0% for all other concentrations (Fig. 5A). In contrast, co-culture in compound **3** resulted in 100% of the parasites being PI positive down to 0.94 µM (Fig. 5B); this result tightly aligns to the motility scores quantified by light microscopy (Fig. 4). Therefore, compounds **1** and **3** are not simply immobilising the juvenile parasites but are, in fact, killing them. To note, control juvenile parasites were also assessed for PI uptake (Fig. 5C). For DMSO treated parasites, as expected with high motility scores, 0% of the parasites scored PI positive. Interestingly, PZQ treated parasites also displayed similar results to that of DMSO treated parasites where 0% of the juvenile worms were PI positive. This is in contradiction to PZQ-treated parasites scoring the lowest (0-1) for motility (Fig. 4) and illustrates that, while this drug decreases motility, it does not kill juvenile stage parasites. Auranofin-treated parasites were all (100%) PI positive.

Evaluation of the indicative cytotoxic effect of these compounds against HepG2 cells was subsequently tested. Compounds **1** - **6** were titrated (from 200 - 1 µM) on HepG2 cells and co-cultivated for 24 hrs. A previous large scale mammalian cytotoxicity study indicated that maximal HepG2 cytotoxicity was observed within the first 24 hrs for 91% of the active compounds [34]; therefore, 24 hrs continuous co-incubation of N-acyl homoserine lactones with HepG2 cells was selected for this study. The racemic compound (**1**) had an EC_50_ of 43.5 µM (Table 2) on HepG2 cells. Evaluation of the individual enantiomers demonstrated that (*R*)- and (*S*)- enantiomers had higher EC_50_ concentrations compared to the parent compound **1** (50.6 µM and >100 µM respectively). Incorporation of the thiolactone (**3**) resulted in greater cytotoxicity (EC_50_ = 18.7µM) with a 2.3 fold increase in comparison to compound **1**.

With the EC_50_ data collected for schistosomula, adult and juvenile worms as well as the CC_50_ for HepG2, the selectivity indices (SI) could be determined for each of the compounds tested (Table 2). For all lifecycle stages tested, the thiolactone analogue (**3**) had the lowest EC_50_ values; however, it also had the poorest CC_50_ values of all the compounds tested, which resulted in some of the lowest SI scores except for juvenile worms where a SI of 37.4 was noted.

Due to limited information regarding potential targets of compound **1**, we next conducted an evaluation of structural similarities to previously published compounds as a first step towards this goal (Fig. 6).

**Figure 6:**
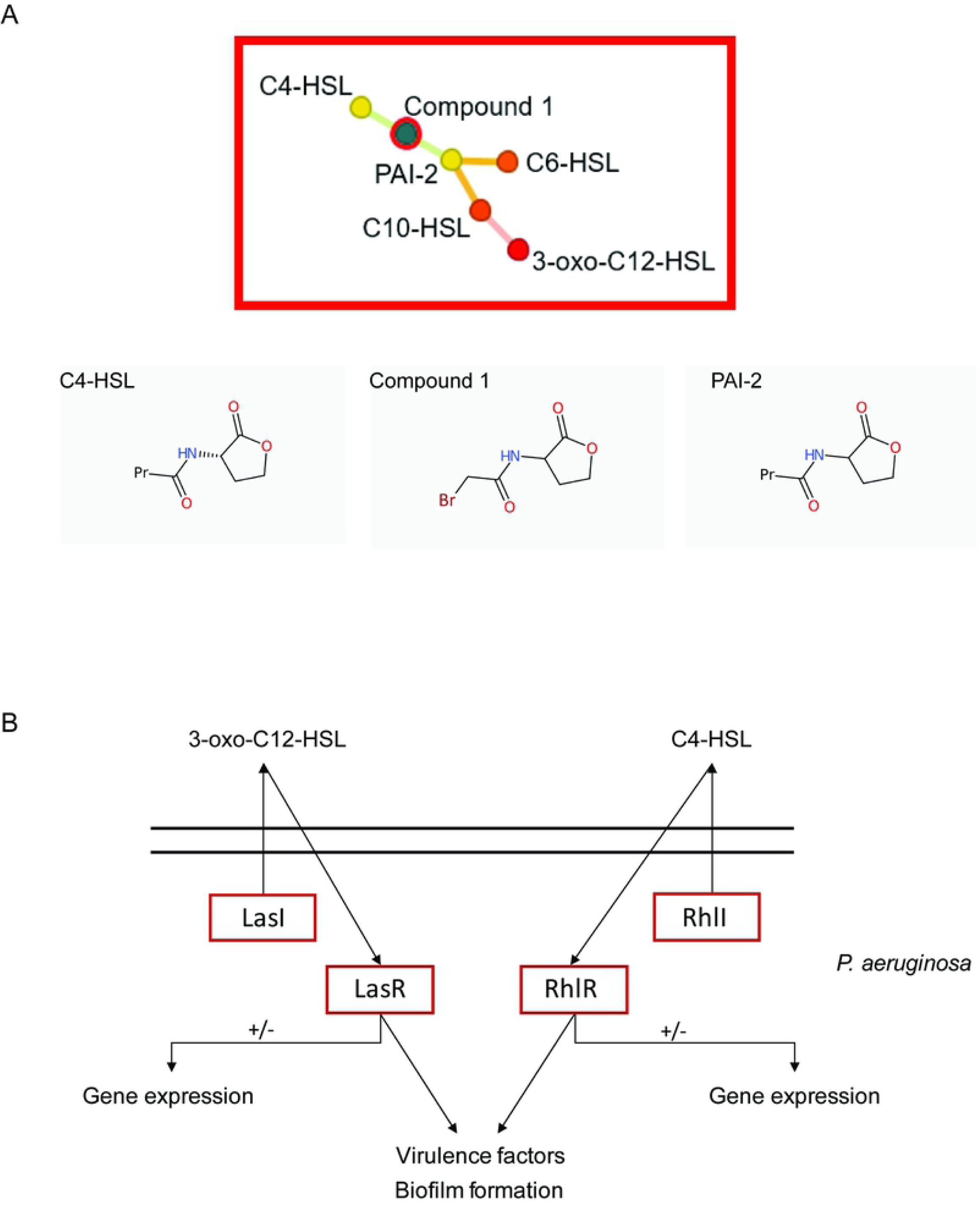
*In silico* approach to identify potential targets of compound 1 within *S. mansoni*. **A)** FragFP output in DataWarrior demonstrates that Compound **1** is structurally related to C4-HSL (formerly called PAI-2 [35]); C4-HSL is an N-acyl homoserines involved in quorum sensing within *P. aeruginosa*. Substitution of the methyl and bromine groups (found in compound **1**) is observed with propyl and stereochemistry modifications. **B)** Pathways in *P. aeruginosa* that utilise both C4- and C12-HSLs (LasI = acyl-homoserine-lactone synthase; LasR = transcriptional activator; RhlR = regulatory protein RhlI = acyl-homoserine-lactone synthase). Highlighted (red rectangles) are those *P. aeruginosa* proteins used in BLASTp analysis of the *S. mansoni* genome (v 7.0).

Using LeadMine and Datawarrior tools [31], compound **1** was found to be structurally similar to that of the N-butanoyl-l-homoserine lactones C4-HSL and PAI-2 (Fig. 6A). Within *P. aeruginosa*, the N-3-oxo-dodecanoyl-l-homoserine lactone C12 and C4 signal through LasR and RhIR to facilitate quorum sensing [36] (Fig. 6B). In addition to this function, both C4 and C12 HSLs regulate gene expression within *P. aeruginosa* as well as within several mammalian host cells [37-39]. Subsequently, we evaluated whether compound **1** may target similar *S. mansoni* orthologues to those used by *P. aeruginosa* in facilitating C4-HSL and C12-HSL signal transduction events. Upon BLATSP analyses of the *S. mansoni* genome, our findings failed to provide convincing evidence for LasR, LasI, RhIR, RHII (*P. aeruginosa)* orthologues (data not shown). This suggests that our N-acyl homoserine analogue (**1**) is operating through differing mechanisms to that seen within bacteria, as has been postulated for mammalian systems [38, 39].

As Compound **1** was structurally similar to N-acyl homoserine lactones, a compound class involved in bacterial quorum sensing, we decided to assess its antimicrobial activity and determine if enantiomer separation or lactone ring substitution would change this activity. Compounds **1, 3, 5** and **6** were thus screened against representative Gram^+^ (*Staphylococcus aureus*) and Gram^-^ (*Escherichia coli*) species at final concentrations of 125, 62.5, 31.25, 15.6, 7.8 and 3.9 mg/L. Minimum inhibitory concentrations (MIC) were determined and represent the minimum concentration associated with no visible bacterial growth. The racemic compound (**1)** showed a MIC of 125 mg/L for both bacteria species, which was a similar value to that obtained for the (*R*)-enantiomer (**5)**. In contrast, the (*S*)-enantiomer (**6**) did not show any activity (> 125 mg/L) on either bacterial species indicating that, similarly to the anthelmintic activity, the (*R*)-enantiomer alone is responsible for the antimicrobial activity (Table 3). When the lactone ring was replaced with a thiolactone substituent (**3**), the activity was considerably improved leading to a MIC of 31.25 mg/L for *S. aureus* (IC_50_ = 25.9 mg/L) and 62.5 mg/L for *E. coli* (IC_50_ = 52.7 mg/L) (Table 3).

**Table 3.** Antibacterial activity (MIC) of the tested compounds and antibacterial selectivity of compound 3 (IC_50_)

| Compound ID | *S. aureus* | *E. coli* | *S. aureus* | *E. coli* | CC<sub>50</sub> | Selectivity Index |
| --- | --- | --- | --- | --- | --- | --- |
|  | MIC |  | IC<sub>50</sub> (μM) |  | (μM)<br>(HepG2) | *S. aureus* /<br>*E.coli* |
| 1 | 125<br>[563.1] | 125<br>[563.1] |  |  |  |  |
| 3 | 31.3<br>[131.2] | 62.5<br>[262.5] | 25.9<br>(23.3 -<br>30.7) | 52.7<br>(42.3 –<br>66.9) | 18.7<br>(14.2 -<br>23.2) | 0.72 / 0.35 |
| 5 | 125<br>[563.1] | 125<br>[563.1] |  |  |  |  |
| 6 | > 125<br>[> 563.1] | > 125<br>[> 563.1] |  |  |  |  |

Minimum inhibitory concentration of compounds expressed as mg/L [μM] against *Staphylococcus aureus* and *Escherichia coli*. IC<sub>50</sub> against *S. aureus* and *E. coli* determined from three independent experiments and expressed as mg/L [μM] with 95% confidence interval (in parentheses). Selectivity indices were calculated from HepG2 CC<sub>50</sub> values.

## Discussion

The drive to identify new anti-infectives is of paramount importance due to the continuous documentation and threat of drug resistant pathogens [40-43]. In particular, our research group has focused on identifying new chemotherapeutic compounds active against *S. mansoni*, one of three major species responsible for human schistosomiasis.

In this study, 14 compounds were initially screened against schistosome parasites. It was evident that compound (**1**) had good activity against a range of lifecycle stages (schistosomula, adult and juvenile worms); therefore, a further 8 analogues (including the separated enantiomers) were generated. Interestingly the (*R*)-enantiomer was more effective when compared to the (*S*)-enantiomer in all of the parasitic assays conducted in this study; the (*R*)-enantiomer alone was also less toxic than parent compound **1** (EC_50_ 50.6µM and 43.5µM respectively). It is not surprising that different activities are associated with enantiomers; indeed PZQ itself is only effective via its (*R*)- enantiomer [44]. For example, Patterson *et al* demonstrated that the (*R*)- enantiomer of PA-824 (a promising antitubercular drug) had greater activity (when compared to the (*S*) enantiomer) against *L. donovani* during *in vivo* studies [45]. Furthermore, Paredes *et al* demonstrated that (*R*)-albendazole sulfoxide had greater anthelmintic activity against *T. solium* when compared to (*S*)-albendazole [46]. Our data, along with these and other parasitic studies [47], provides clear evidence that compound chirality represents an important consideration for drug discovery progression and putative target identification.

Upon further exploration of structural activity relationships, we found that elongation of the N-acyl chain led to decreased activity (**1** vs **2**; **3** vs **4**), while substitution of the lactone ring with a thiolactone improved activity (**1** vs **3, 2** vs **4**) and led to the most active compound **3**. It is reasonable to speculate that should **3** be available as the pure (*R*)-enantiomer, improvements in both anti-schistosomal potency and host cell cytotoxicity would be found (similar to those observed for compound **1** versus enantiomer pure (*R*) - **5** and (*S*) - **6**). A reduction in schistosome fecundity was noted in cultures co-incubated with **3** vs **4;** therefore, it could be argued that the addition of the methyl group in position 2 to the thiolactone analogue reduces parasite fecundity.

Comparative structural analysis revealed that compound **1**, having been previously characterised as an intermediary product of one-step procedures for producing Gram^-^ N-acyl homoserines [48, 49], was similar to that of quorum sensing N-acyl homoserines. In recent years, quorum sensing has become a focus of development as a target for new anti-infective treatments [50]. Quorum sensing is a method by which both eukaryotic and prokaryotic cells regulate gene expression in response to fluctuations in cell population densities. Of relevance to parasites, *Trypanosoma brucei* is known to regulate its surrounding population by an analogous quorum sensing mechanism. In order to evade the host immunity response, trypanosomes adopt a slender morphology to proliferate, and in order to enter their transmission stage will transform into a stumpy form in which they stop proliferating [51]. This differentiation between two morphologically and molecular lifecycle stages is dependent on parasite density, and therefore can be seen as a type of quorum sensing [52, 53]. Much work has gone into identifying the mechanisms responsible for this density sensing to provide potential quorum sensing drug targets in trypanosomes [53]. Quorum sensing in schistosomes (or other metazoan endoparasites), to regulate lifecycle transitions, has not yet been identified.

With the identification of our parent compound having structural similarities to that of quorum-sensing N-acyl homoserines, we decided to evaluate compounds **1** and **3** (the most effective compound against *S. mansoni*), as well as the (*R*)- and (*S*)- enantiomers (**5** and **6** respectively) against both Gram^+^ and Gram^-^ bacteria to assess their potential as antimicrobials. Our findings in the bacteria screens were consistent with the *S. mansoni* data in that **3** was the most potent N-acyl homoserine against Gram^+^ bacteria and that the (*R*)- (**5**) was more effective than the (*S*)- (**6**) enantiomer. Furthermore, the thiolactone (**3**) had a 4- and 2- fold improvement in activity against *S. aureus* and *E. coli* respectively, compared to compound **1**. It is interesting that N-acyl homoserines should have greater effect against a Gram^+^ bacteria, which in natural environments do not utilise these compounds as part of their quorum sensing regulation [54]. An explanation for increased inhibition of *S. aureus* growth when treated with compound **3** may be due to cross-inhibition between bacteria species. It has previously been demonstrated that bacteria undergo strategies to reduce the efficacy of quorum sensing used by competitor bacterial species and thus increase the likelihood of their own survival [54, 55]. The role of N-acyl homoserines in this process of cell to cell bacterial communication has been clearly demonstrated for *S. aureus* and *P. aeruginosa* [56]. Here, N-acyl homoserines (e.g. 3-oxo-C_12_-HSL) produced by Gram^-^ *P. aeruginosa* inhibit growth and virulence factor production of Gram^+^ *S. aureus*. When co-inhabiting infected tissues, this N-acyl homoserine-mediated strategy of cell communication provides *P. aeruginosa* with a competitive advantage over *S. aureus*. Other studies have also demonstrated a role for N-acyl homoserines in mediating pathogen population densities; for example, *Candida albicans* filamentation can be suppressed by 3-oxo-C_12_-HSL (homoserine lactone) produced by *P. aeruginosa* [57]. It is, therefore, likely that our synthetic N-acyl homoserine analogues (especially thiolactone containing compound **3**) inhibit *S. aureus* growth in a similar manner as the above examples or those additional ones found in the literature illustrate (54, 55). For example, McInnis *et al* have previously designed and validated a library of 21 thiolactone analogues that were either naturally occurring or non-native N-acyl homoserines and evaluated their effect as agonists and antagonists of LuxR-type quorum sensing receptors in *P*.*aeruginosa* (LasR), *V. fischeri* (LuxR), and *A*.*tumefaciens* (TraR). They were able to demonstrate the improved potency of the thiolactone analogues in comparison to the N-acyl homoserine parent compound (similar to our findings here) [58].

In *P. aeruginosa*, its N-acyl homoserines primarily function through the transcription factors LasR and RhIR, but similar orthologues were not identified in the *S. mansoni* genome. However, it has also been shown that bacterial N-acyl homoserines mediate cross-Kingdom communication by directly interacting with membrane lipids; this mechanism of action may be responsible for the anti-schistosomal effects observed in our studies (although detailed microscopy studies would need to confirm this hypothesis). For example, long chain N-acyl homoserines (e.g. 3-oxo-C_12_-HSL) can insert into cholesterol-containing microdomains leading to changes in dipole potential, integral protein re-organisation and subsequent activation of signal transduction cascades [37, 59]. The intra-mammalian schistosome is covered by two tightly opposed lipid bilayers including an inner plasma membrane and an outer membranocalyx [60-62]. The membranocalyx has the highest density of intramembranous particles [63] comprised of nutrient transporters, transmembrane proteins and other gene products of currently unknown function [64-66]. Also, it is rich in neutral lipids, cholesterol and phospholipids [67-70]. It is possible that the compounds analysed in this study may bind to the cholesterol rich regions within this membranocalyx layer, which subsequently causes conformational changes to the transmembrane proteins affecting their downstream signalling cascades. This could ultimately result in the phenotypes observed in our studies.

To the best of our knowledge, this is the first study to demonstrate that synthetic bacterial N-acyl homoserines analogues are active against *S. mansoni*. This is a new avenue of investigation as these compounds represent novel starting points for future anthelmintic drug discovery studies. We have also demonstrated the importance of enantiomer contribution, in line with previous findings, and that some molecules display cross-pathogen activity. When taken together, these findings could inform the development of new, broad-acting anti-infectives.

## Acknowledgments

We thank Ms Julie Hirst and all of Prof. Karl F. Hoffmann’s laboratory for help in maintaining the *S. mansoni* life cycle. We also thank Prof. Luis Muir, Aberystwyth University, for the use of his laboratory for conducting the bacterial screens. We also thank Noel O’Boyle (NextMove Software Ltd.) for kindly running LeadMine on PubMed abstracts for this study. For the mass spectrometry data, we would like to thank the EPSRC National Mass Spectrometry Service Centre in Swansea.

## Supporting Information

**S1 Protocol: Detailed materials and methods for chemical synthesis of compounds 1 – 9**.

**S1 Table: Structural details of all compounds screened against *S. mansoni* in this study**. The first 14 compounds represent the initial compounds screened. The following 8 are compound **1** analogues.

**S1 Figure: Initial schistosomula screens of the 14 compounds and their intermediate analogues**. A total of 120 mechanically transformed schistosomula were co-cultured with each compound, titrated at doses between 10 and 0.625 µM. Test plates were incubated at 37°C for 72 hrs in an atmosphere containing 5% CO_2._ At 72hrs, schistosomula were scored using the Roboworm platform for both motility and phenotype as previously described [17, 25]. **A)** Of the 14 compounds evaluated, two fell within the hit region. Z’ scores for motility and phenotype were 0.40079 and 0.56846 respectively. **B)** Further titration of the two hit compounds resulted in good compound effect being observed for compound **1** down to a concentration of 1.25 µM. Z’ scores for motility and phenotype were 0.37864 and 0.47882 respectively. **\***Average EC_50_ value across motility and phenotype is presented for each compound.

## References

1. Organization WH. Working to overcome the global impact of neglected tropical diseases: first WHO report on neglected tropical diseases. Geneva: World Health Organization; 2010.

2. Hotez PJ, Alvarado M, Basáñez M-G, Bolliger I, Bourne R, Boussinesq M, et al. The global burden of disease study 2010: interpretation and implications for the neglected tropical diseases. PLoS neglected tropical diseases. 2014;8(7):e2865.

3. Frieri M, Kumar K, Boutin A. Antibiotic resistance. Journal of infection and public health. 2017;10(4):369–78.

4. Morschhäuser J. Regulation of multidrug resistance in pathogenic fungi. Fungal Genetics and Biology. 2010;47(2):94–106.

5. Chevalier FD, L. Clec’h W, McDew-White M, Menon V, Guzman MA, Holloway SP, et al. Oxamniquine resistance alleles are widespread in Old World Schistosoma mansoni and predate drug deployment. PLoS pathogens. 2019;15(10).

6. Patrick J, Anderson F, Wilson K, McCormick I, Skuce P, O’Roarke J. Triclabendazole-resistant liver fluke: issues and strategies. Livestock. 2018;23(Sup5):4–14.

7. Noedl H, Se Y, Schaecher K, Smith BL, Socheat D, Fukuda MM. Evidence of artemisinin-resistant malaria in western Cambodia. New England Journal of Medicine. 2008;359(24):2619–20.

8. Lira R, Sundar S, Makharia A, Kenney R, Gam A, Saraiva E, et al. Evidence that the high incidence of treatment failures in Indian kala-azar is due to the emergence of antimony-resistant strains of Leishmania donovani. The Journal of infectious diseases. 1999;180(2):564–7.

9. Shlaes DM, Projan SJ, Edwards J. Antibiotic discovery: state of the state. ASM News-American Society for Microbiology. 2004;70(6):275–81.

10. Bush K. Why it is important to continue antibacterial drug discovery. ASM News-American Society for Microbiology. 2004;70(6):282–7.

11. Projan SJ. Why is big Pharma getting out of antibacterial drug discovery? Current opinion in microbiology. 2003;6(5):427–30.

12. El Ridi RA, Tallima HA-M. Novel therapeutic and prevention approaches for schistosomiasis. Journal of Advanced Research. 2013;4(5):467–78.

13. Colley D, Secor W. Immunology of human schistosomiasis. Parasite immunology. 2014;36(8):347–57.

14. Vale N, Gouveia MJ, Rinaldi G, Brindley PJ, Gärtner F, da Costa JMC. Praziquantel for schistosomiasis: single-drug metabolism revisited, mode of action, and resistance. Antimicrobial agents and chemotherapy. 2017;61(5):e02582–16.

15. Cioli D, Pica-Mattoccia L, Basso A, Guidi A. Schistosomiasis control: praziquantel forever? Molecular and biochemical parasitology. 2014;195(1):23–9.

16. Edwards J, Brown M, Peak E, Bartholomew B, Nash RJ, Hoffmann KF. The diterpenoid 7-keto-sempervirol, derived from Lycium chinense, displays anthelmintic activity against both Schistosoma mansoni and Fasciola hepatica. PLoS neglected tropical diseases. 2015;9(3):e0003604.

17. Crusco A, Bordoni C, Chakroborty A, Whatley KC, Whiteland H, Westwell AD, et al. Design, synthesis and anthelmintic activity of 7-keto-sempervirol analogues. European journal of medicinal chemistry. 2018;152:87–100.

18. Whiteland HL, Chakroborty A, Forde-Thomas JE, Crusco A, Cookson A, Hollinshead J, et al. An Abies procera-derived tetracyclic triterpene containing a steroid-like nucleus core and a lactone side chain attenuates in vitro survival of both Fasciola hepatica and Schistosoma mansoni. International Journal for Parasitology: Drugs and Drug Resistance. 2018;8(3):465–74.

19. Whatley K, Padalino G, Whiteland H, Geyer K, Hulme B, Chalmers I, et al. The repositioning of epigenetic probes/inhibitors identifies new anti-schistosomal lead compounds and chemotherapeutic targets. BioRxiv. 2019:729814.

20. Padalino G, Ferla S, Brancale A, Chalmers IW, Hoffmann KF. Combining bioinformatics, cheminformatics, functional genomics and whole organism approaches for identifying epigenetic drug targets in Schistosoma mansoni. International Journal for Parasitology: Drugs and Drug Resistance. 2018;8(3):559–70.

21. Crusco A, Baptista R, Bhowmick S, Beckmann M, Mur LA, Westwell AD, et al. The anti-mycobacterial activity of a diterpenoid-like molecule operates through nitrogen and amino acid starvation. Frontiers in Microbiology. 2019;10:1444.

22. Williams P. Quorum sensing, communication and cross-kingdom signalling in the bacterial world. Microbiology. 2007;153(12):3923–38.

23. Bickle Q, Doenhoff M. Comparison of the live vaccine potential of different geographic isolates of Schistosoma mansoni. Journal of helminthology. 1987;61(3):191–5.

24. Colley DG, Wikel SK. Schistosoma mansoni: simplified method for the production of schistosomules. Exp Parasitol. 1974;35(1):44–51.

25. Crusco A, Bordoni C, Chakroborty A, Whatley KCL, Whiteland H, Westwell AD, et al. Design, synthesis and anthelmintic activity of 7-keto-sempervirol analogues. Eur J Med Chem. 2018;152:87–100.

26. Ramirez B, Bickle Q, Yousif F, Fakorede F, Mouries MA, Nwaka S. Schistosomes: challenges in compound screening. Expert Opin Drug Discov. 2007;2(1):S53–61.

27. Peak E, Chalmers IW, Hoffmann KF. Development and validation of a quantitative, high-throughput, fluorescent-based bioassay to detect schistosoma viability. PLoS neglected tropical diseases. 2010;4(7):e759.

28. Nur EAM, Yousaf M, Ahmed S, Al-Sheddi ES, Parveen I, Fazakerley DM, et al. Neoclerodane Diterpenoids from Reehal Fatima, Teucrium yemense. J Nat Prod. 2017;80(6):1900–8.

29. Microbiology ECfASTotESoC, Diseases I. Determination of minimum inhibitory concentrations (MICs) of antibacterial agents by broth dilution. Clinical Microbiology and Infection. 2003;9(8):ix–xv.

30. Lowe DM, Sayle RA. LeadMine: a grammar and dictionary driven approach to entity recognition. Journal of cheminformatics. 2015;7(1):S5.

31. Sander T, Freyss J, von Korff M, Rufener C. DataWarrior: an open-source program for chemistry aware data visualization and analysis. Journal of chemical information and modeling. 2015;55(2):460–73.

32. Howe KL, Bolt BJ, Cain S, Chan J, Chen WJ, Davis P, et al. WormBase 2016: expanding to enable helminth genomic research. Nucleic acids research. 2016;44(D1):D774–D80.

33. Gray DJ, Ross AG, Li Y-S, McManus DP. Diagnosis and management of schistosomiasis. Bmj. 2011;342:d2651.

34. Hsieh J-H, Huang R, Lin J-A, Sedykh A, Zhao J, Tice RR, et al. Real-time cell toxicity profiling of Tox21 10K compounds reveals cytotoxicity dependent toxicity pathway linkage. PloS one. 2017;12(5):e0177902.

35. Pearson JP, Van Delden C, Iglewski BH. Active efflux and diffusion are involved in transport of Pseudomonas aeruginosa cell-to-cell signals. Journal of bacteriology. 1999;181(4):1203–10.

36. Pesci EC, Pearson JP, Seed PC, Iglewski BH. Regulation of las and rhl quorum sensing in Pseudomonas aeruginosa. Journal of bacteriology. 1997;179(10):3127–32.

37. Davis BM, Jensen R, Williams P, O’Shea P. The interaction of N-acylhomoserine lactone quorum sensing signaling molecules with biological membranes: implications for inter-kingdom signaling. PloS one. 2010;5(10):e13522.

38. Kravchenko VV, Kaufmann GF, Mathison JC, Scott DA, Katz AZ, Wood MR, et al. N-(3-oxo-acyl) homoserine lactones signal cell activation through a mechanism distinct from the canonical pathogen-associated molecular pattern recognition receptor pathways. Journal of Biological Chemistry. 2006;281(39):28822–30.

39. Hooi DS, Bycroft BW, Chhabra SR, Williams P, Pritchard DI. Differential immune modulatory activity of Pseudomonas aeruginosa quorum-sensing signal molecules. Infection and immunity. 2004;72(11):6463–70.

40. Tacconelli E, Carrara E, Savoldi A, Harbarth S, Mendelson M, Monnet DL, et al. Discovery, research, and development of new antibiotics: the WHO priority list of antibiotic-resistant bacteria and tuberculosis. The Lancet Infectious Diseases. 2018;18(3):318–27.

41. Nikaido H. Multidrug resistance in bacteria. Annual review of biochemistry. 2009;78:119–46.

42. Donia M, Hamann MT. Marine natural products and their potential applications as anti-infective agents. The Lancet infectious diseases. 2003;3(6):338–48.

43. Doenhoff MJ, Kusel JR, Coles GC, Cioli D. Resistance of Schistosoma mansoni to praziquantel: is there a problem? Transactions of the Royal Society of Tropical Medicine and Hygiene. 2002;96(5):465–9.

44. Meister I, Ingram-Sieber K, Cowan N, Todd M, Robertson MN, Meli C, et al. Activity of praziquantel enantiomers and main metabolites against Schistosoma mansoni. Antimicrobial agents and chemotherapy. 2014;58(9):5466–72.

45. Patterson S, Wyllie S, Stojanovski L, Perry MR, Simeons FR, Norval S, et al. The R enantiomer of the antitubercular drug PA-824 as a potential oral treatment for visceral leishmaniasis. Antimicrobial agents and chemotherapy. 2013;57(10):4699–706.

46. Paredes A, de Campos Lourenço T, Marzal M, Rivera A, Dorny P, Mahanty S, et al. In vitro analysis of albendazole sulfoxide enantiomers shows that (+)-(R)-albendazole sulfoxide is the active enantiomer against Taenia solium. Antimicrobial agents and chemotherapy. 2013;57(2):944–9.

47. Nanayakkara ND, Ager AL, Bartlett MS, Yardley V, Croft SL, Khan IA, et al. Antiparasitic activities and toxicities of individual enantiomers of the 8-aminoquinoline 8-[(4-amino-1-methylbutyl) amino]-6-methoxy-4-methyl-5-[3, 4-dichlorophenoxy] quinoline succinate. Antimicrobial agents and chemotherapy. 2008;52(6):2130–7.

48. Persson T, Hansen TH, Rasmussen TB, Skindersø ME, Givskov M, Nielsen J. Rational design and synthesis of new quorum-sensing inhibitors derived from acylated homoserine lactones and natural products from garlic. Organic & biomolecular chemistry. 2005;3(2):253–62.

49. Stacy DM, Le Quement ST, Hansen CL, Clausen JW, Tolker-Nielsen T, Brummond JW, et al. Synthesis and biological evaluation of triazole-containing N-acyl homoserine lactones as quorum sensing modulators. Organic & biomolecular chemistry. 2013;11(6):938–54.

50. Kaufmann GF, Park J, Janda KD. Bacterial quorum sensing: a new target for anti-infective immunotherapy. Expert opinion on biological therapy. 2008;8(6):719–24.

51. Vickerman K. Developmental cycles and biology of pathogenic trypanosomes. British medical bulletin. 1985;41(2):105–14.

52. Vassella E, Reuner B, Yutzy B, Boshart M. Differentiation of African trypanosomes is controlled by a density sensing mechanism which signals cell cycle arrest via the cAMP pathway. Journal of cell science. 1997;110(21):2661–71.

53. Mony BM, MacGregor P, Ivens A, Rojas F, Cowton A, Young J, et al. Genome-wide dissection of the quorum sensing signalling pathway in Trypanosoma brucei. Nature. 2014;505(7485):681.

54. Bassler BL. Small talk: cell-to-cell communication in bacteria. Cell. 2002;109(4):421–4.

55. Taga ME, Bassler BL. Chemical communication among bacteria. Proceedings of the National Academy of Sciences. 2003;100(Suppl 2):14549–54.

56. Qazi S, Middleton B, Muharram SH, Cockayne A, Hill P, O’Shea P, et al. N-acylhomoserine lactones antagonize virulence gene expression and quorum sensing in Staphylococcus aureus. Infection and immunity. 2006;74(2):910–9.

57. Hogan DA, Vik Å, Kolter R. A Pseudomonas aeruginosa quorum-sensing molecule influences Candida albicans morphology. Molecular microbiology. 2004;54(5):1212–23.

58. McInnis CE, Blackwell HE. Thiolactone modulators of quorum sensing revealed through library design and screening. Bioorganic & medicinal chemistry. 2011;19(16):4820–8.

59. Song D, Meng J, Cheng J, Fan Z, Chen P, Ruan H, et al. Pseudomonas aeruginosa quorum-sensing metabolite induces host immune cell death through cell surface lipid domain dissolution. Nature microbiology. 2019;4(1):97–111.

60. Hockley DJ, McLaren DJ. Schistosoma mansoni: changes in the outer membrane of the tegument during development from cercaria to adult worm. International journal for parasitology. 1973;3(1):13–20.

61. McLaren DJ, Hockley DJ. Blood flukes have a double outer membrane. Nature. 1977;269(5624):147–9.

62. Skelly PJ, Wilson RA. Making sense of the schistosome surface. Advances in parasitology. 2006;63:185–284.

63. Hockley D, McLaren DJ, Ward BJ, Nermut M. A freeze-fracture study of the tegumental membrane of Schistosoma mansoni (Platyhelminthes: Trematoda). Tissue and Cell. 1975;7(3):485–96.

64. van Balkom BW, van Gestel RA, Brouwers JF, Krijgsveld J, Tielens AG, Heck AJ, et al. Mass Spectrometric Analysis of the Schistosoma m ansoni Tegumental Sub-proteome. Journal of proteome research. 2005;4(3):958–66.

65. Braschi S, Borges WC, Wilson RA. Proteomic analysis of the shistosome tegument and its surface membranes. Memorias do Instituto Oswaldo Cruz. 2006;101:205–12.

66. Castro-Borges W, Simpson DM, Dowle A, Curwen RS, Thomas-Oates J, Beynon RJ, et al. Abundance of tegument surface proteins in the human blood fluke Schistosoma mansoni determined by QconCAT proteomics. Journal of Proteomics. 2011;74(9):1519–33.

67. Retra K, deWalick S, Schmitz M, Yazdanbakhsh M, Tielens AG, Brouwers JF, et al. The tegumental surface membranes of Schistosoma mansoni are enriched in parasite-specific phospholipid species. International journal for parasitology. 2015;45(9-10):629–36.

68. Tallima H, Hamada M, El Ridi R. Evaluation of cholesterol content and impact on antigen exposure in the outer lipid bilayer of adult schistosomes. Parasitology. 2007;134(12):1775–83.

69. Rogers MV, McLaren DJ. Analysis of total and surface membrane lipids of Schistosoma mansoni. Molecular and biochemical parasitology. 1987;22(2-3):273–88.

70. Migliardo F, Tallima H, El Ridi R. Is There a Sphingomyelin-Based Hydrogen Bond Barrier at the Mammalian Host–Schistosome Parasite Interface? Cell biochemistry and biophysics. 2014;68(2):359–67.

